# No molecular evidence for Muller’s ratchet in mitochondrial genomes

**DOI:** 10.64898/2025.12.16.694700

**Authors:** Yu K. Mo, Sarthak R. Mishra, Yadira Peña-Garcia, Matthew W. Hahn

## Abstract

Muller’s ratchet predicts that non-recombining genomes can accumulate deleterious mutations, though molecular evidence for it is rare. Previous studies have tried to detect ratchet-like behavior in mitochondria (mtDNA), contrasting the rate of nucleotide substitutions in mitochondrial transfer RNA (tRNA) genes with those in the nuclear genome. However, these studies relied on small datasets that could not control for the higher mutation rate in animal mtDNA. In this study, we re-evaluate evidence for Muller’s ratchet with well-annotated mitochondrial and nuclear genomes from three species pairs. Across all three comparisons, mitochondrial tRNA (and protein-coding) genes are evolving under strong constraint, with *dt*/*dS* equal to 0.12 in primates, 0.07 in birds, and 0.04 in fruit flies. Compared with nuclear *dt*/*dS*, only the fruit fly comparison showed slightly lower constraint in mtDNA. Our results therefore provide evidence for strong, efficacious selection in animal mtDNA, with no ratchet-like accumulation of slightly deleterious mutations.

## Introduction

Muller’s ratchet predicts that genomes without recombination will accumulate deleterious mutations (Muller 1964; Felsenstein 1974). In the absence of back-mutation, and under a specific set of assumptions about the strength and type of selection, quantitative predictions can be made about the speed at which genomes affected by the ratchet will accumulate slightly deleterious mutations (Haigh 1978; Charlesworth and Charlesworth 1997; Gordo and Charlesworth 2000). However, it has been hard to test specific predictions about the ratchet, at least in part because genomes that have recently stopped recombining have likely also experienced large shifts in their selective environments (Chao 1990; Bachtrog and Gordo 2004) or in their molecular machinery (Moran 1996; Itoh et al. 2002).

A genomic compartment long thought to be subject to Muller’s ratchet is the mitochondrion (Gabriel et al. 1993; Neiman and Taylor 2009). MtDNA is largely non-recombining and usually uniparentally inherited. In animals, mtDNA also has a higher mutation rate compared with the nuclear genome (Pesole et al. 1999; Ballard and Whitlock 2004; Haag-Liautard et al. 2008); in plants, the nuclear genome usually has the higher rate (but see Sloan et al. 2012). One straightforward prediction of Muller’s ratchet is that the mtDNA genome will show a “wider selective sieve” than the nuclear genome (Lynch 1997), allowing the fixation of slightly deleterious mutations. However, when this has been tested with protein-coding genes, the mtDNA has shown either equivalent or stronger signals of purifying selection (Popadin et al. 2013; Cooper et al. 2015).

One problem with the comparison of mtDNA and nuclear protein-coding genes is that the functions of mtDNA genes are highly specialized and extremely important to organismal function (Ballard and Whitlock 2004); comparing genes with very different functions between compartments would not be a good test of Muller’s ratchet. A set of genes that carry out the same function between the mtDNA and nuclear genomes is tRNAs. In an influential set of papers, Lynch (1996; 1997) compared the rate of nucleotide substitution in tRNAs between the mtDNA and nuclear genomes. He found that the rate in mtDNA genes was substantially higher than in nuclear genes, suggesting evidence for Muller’s ratchet.

However, the results from Lynch (1996; 1997) come with several caveats. First, these studies relied on relatively small datasets, with small numbers of tRNA genes in both genomic compartments. Because of the limited numbers of sequenced genes at the time, different tRNAs were compared between different species, across both mtDNA genes and across nuclear genes; there was no way to consistently compare tRNA genes across compartments between the same set of species. Additionally, due to the lack of sequenced genomes, orthology assignments between (especially) nuclear tRNAs was difficult, as there are many copies of these genes in nuclear genomes. Finally, although it was known that mtDNA has a higher mutation rate, there was no way to control for this difference in the comparisons, except indirectly using divergence times assumed for a phylogenetic tree.

Here, we aimed to re-evaluate evidence for Muller’s ratchet in animal mtDNA with a larger dataset and complete genomes. We use well-annotated genomes to examine tRNA and protein-coding genes from both mitochondrial and nuclear genomes, thereby increasing the number of genes available for comparison. We controlled for differences in mutation rate between compartments by using synonymous substitutions estimated from protein-coding sequences in the same compartment. To avoid problems with possibly discordant gene trees—and to ensure high-quality annotations and orthology assignments—we focused on three pairwise comparisons: human–rhesus macaque (primates), chicken–turkey (birds), and *D. melanogaster*–*D. simulans* (fruit flies).

## Results

After careful filtering, for all three comparisons we retained almost all mtDNA loci (Supplementary Table 1), both tRNA genes and coding sequences (“CDS”). Only three genes total were removed from the mtDNA datasets (Materials and Methods). The numbers of nuclear tRNA and CDS were both higher for all comparisons—hundreds of tRNA genes and thousands of CDS alignments (Supplementary Table 1)—though a much larger fraction of such loci were filtered out due to orthology, alignment, or other issues (see Materials and Methods).

As expected due to the higher mutation rate in mtDNA, the number of synonymous substitutions between species (as measured by the number of synonymous substitutions per-site in CDS), are much higher in mtDNA genes (*dS*_mt_) than nuclear genes (*dS*_n_). The mean *dS*_mt_ is approximately 17.7× higher than *dS*_n_ in primates, 9.7× higher in birds, and 4.3× higher in fruit flies (Table 1; Supplementary Figure 1a–c). Similarly, nonsynonymous substitutions are higher in mtDNA genes than in the nucleus for primates and birds, though this pattern is slightly reversed in fruit flies (Supplementary Figure 1f; Supplementary Table 2).

**Table 1:**
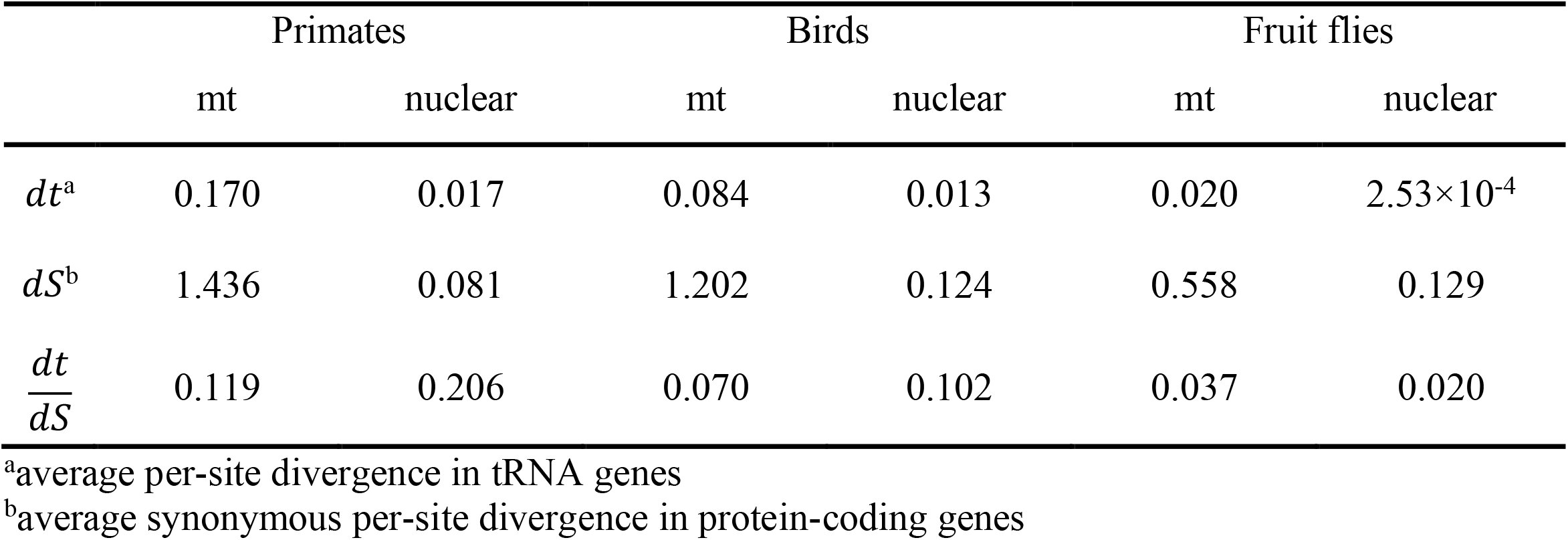
Divergence of tRNA genes and synonymous substitutions for mt and nuclear genes.

A common and powerful way to measure the strength and type of selection is to normalize nonsynonymous substitutions by synonymous substitutions, resulting in the *dN*/*dS* ratio (Kimura 1977; Hurst 2002). If there are no advantageous substitutions, then 1-*dN*/*dS* represents the fraction of all nonsynonymous substitutions that are removed by selection; if there are advantageous substitutions, it is a lower bound on this quantity (Hahn 2018, chapter 7). Lower values therefore represent stronger purifying selection. In all three comparisons, *dN/dS* among mtDNA protein-coding genes was very low, ranging from 0.017 in fruit flies to 0.093 in primates (Supplementary Table 2). Comparison of *dN/dS* between mtDNA and nuclear compartments revealed lower values in the mitochondria for primates and birds, and only slightly higher in fruit flies (Figure 1a–c; Supplementary Table 2). For fruit flies, one mtDNA CDS (*ND4*) had a higher *dN/dS* than the others (=1.84). If we exclude this gene, the mean mitochondrial *dN/dS* drops from 0.173 to 0.0218, and the mtDNA is once again lower than the nuclear genome. Regardless, these patterns indicate strong purifying selection on mtDNA protein-coding genes despite elevated mutation rates.

Similarly to using *dN/dS* in protein-coding genes, we can normalize divergence in tRNA genes (*dt*) by the average synonymous divergence in the appropriate compartment. That is, while we cannot measure *dS* for individual tRNA genes, we can (for instance) divide *dt* in each mtDNA tRNA by the average *dS* among mtDNA CDS. If the mutation rate is the same or similar among genes within the same compartment, *dt/dS* will be proportional to the strength of purifying selection on tRNAs (see Discussion).

As with *dS*_mt_ and *dN*_mt_, divergence among mtDNA tRNA genes (*dt*_mt_) is consistently higher than among nuclear tRNA genes (*dt*_n_), for which there are a number of identical pairs in all three species comparisons (Table 1; Supplementary Figure 1g–i). Because there are hundreds of nuclear tRNA genes, often residing in arrays, assigning orthology among them could be difficult. To ensure that our estimates of *dt*_n_ were not inflated due to comparisons among non-orthologous loci, we compared our nuclear tRNA results between humans and macaques to those found in Thornlow et al. (2018). Our results were highly similar to those in Thornlow et al., both in terms of the distribution of *dt* values (Supplementary Figure 2a) and in terms of the number of orthologous tRNA genes found on each chromosome (Supplementary Figure 2b). These comparisons provide confidence in our methods for identifying orthologs and measuring divergence in tRNAs.

When we normalize tRNA divergence by synonymous divergence, the ratio *dt*_mt_*/dS*_mt_ is very low, ranging from 0.037 in fruit flies to 0.119 in primates (Table 1; Figure 1d–f). These values are lower than, or as low as (depending on the species pair), the corresponding values of *dN/dS* in protein-coding mtDNA genes, emphasizing the selective importance of tRNA genes. Compared to values among nuclear tRNA genes, *dt*_mt_*/dS*_mt_ is again lower in primates and birds, but slightly higher in fruit flies (Table 1; Figure 1d–f). As the level of constraint on even the fruit fly mitochondrial tRNAs implies that >96% of mutations at these loci have been eliminated by selection, these results demonstrate strong selection on mtDNA.

**Figure 1.**
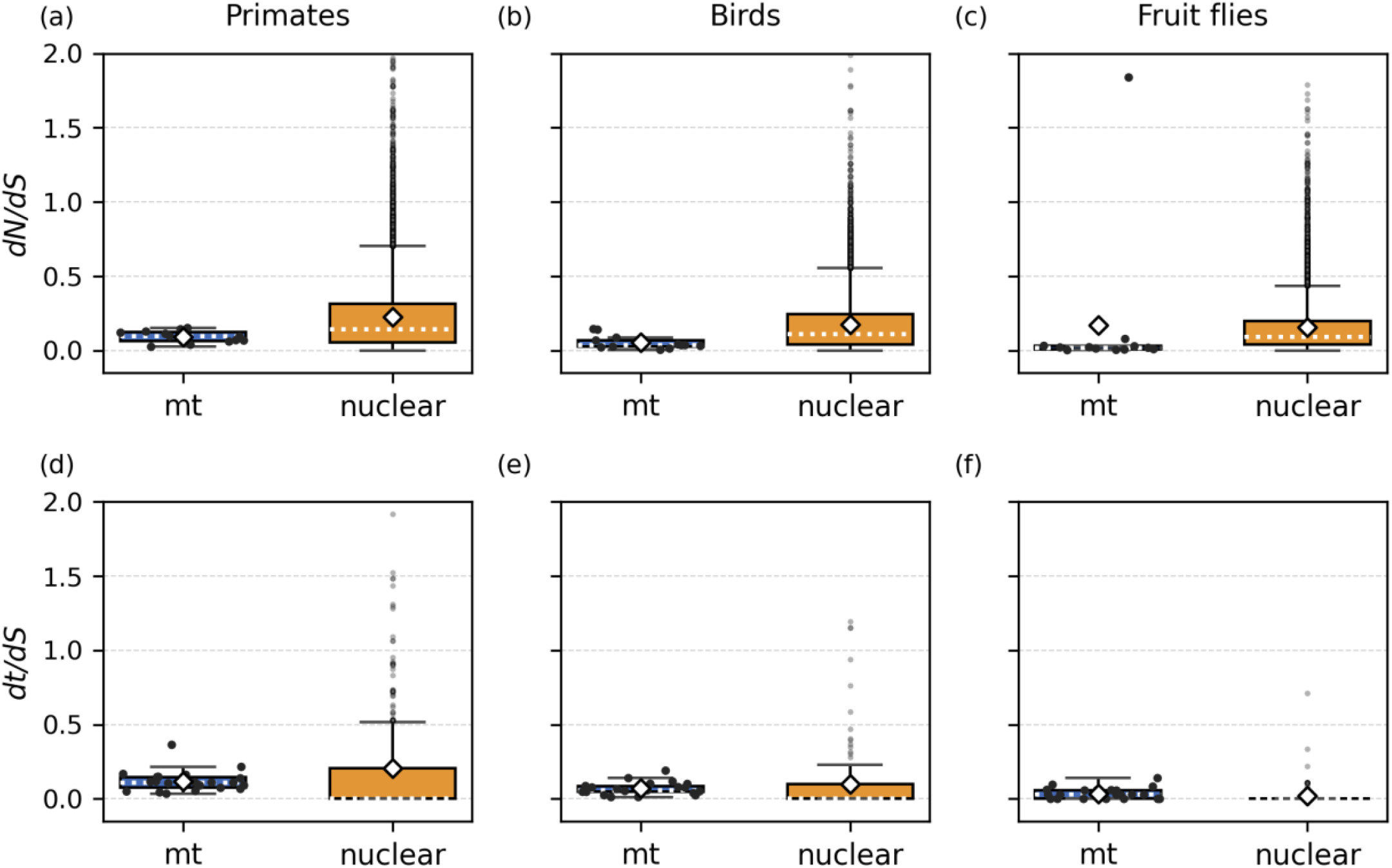
Distributions of *dN/dS* and *dt/dS* for mitochondrial (mt) and nuclear genes across three species pairs. (a)–(c) *dN/dS* for protein-coding genes in primates, birds, and fruit flies, respectively. Panels (d)–(f) *dt/dS* for tRNAs in the same comparisons. White diamonds indicate mean values, and dashed lines are median values. Black dots represent values for individual mtDNA genes; gray dots are nuclear outliers, whose values are greater than Q3 + 1.5×IQR. The y-axes in panels (d) and (e) are truncated, omitting one and four nuclear points with *dt/dS* > 2 (these points were included in all calculations).

## Discussion

Muller’s ratchet has been the subject of intense scrutiny by evolutionary biologists for over 50 years (cf. Felsenstein 1974). However, while the theoretical basis and properties of the ratchet are now well-understood, empirical tests of its action are much rarer. Mitochondria seem to be the perfect setting for the ratchet to operate, as they generally lack recombination and have small(er) population sizes. However, early attempts to demonstrate a relative lack of constraint in mtDNA were limited by the availability of both mitochondrial and nuclear genomes (Lynch 1996, 1997; Lynch and Blanchard 1998). Here, using whole genomes, we find that both protein-coding genes and tRNA genes are under very strong constraint in mtDNA, similar to their nuclear counterparts (see Popadin et al. 2013; Cooper et al. 2015 for similar results on protein-coding genes).

To paraphrase Kondrashov (1995): why has the mitochondrion not died 100 times over? That is, how could we possibly find no evidence for Muller’s ratchet? The simple answer is that for the ratchet to work, a long list of assumptions must be true. Loewe (2006) reviews these assumptions in detail, but here we discuss only a few that may be most relevant. First, the ratchet assumes no recombination and no back-mutation. We now know that homologous recombination in mtDNA can occur (Klucnika et al. 2023), though it can only act to remove deleterious alleles in heteroplasmic individuals; it may therefore be effectively quite rare. Back-mutations may be much more common, especially after early deleterious mutations have fixed (Charlesworth 2012). Second, and more subtly, the ratchet assumes a specific distribution of mutation rates and fitness effects (Haigh 1978; Gessler 1995) and no epistasis (Kondrashov 1994; but see Butcher 1995). It is not clear whether mtDNA matches these assumptions. Third, the ratchet assumes there is no transfer of material into the non-recombining genome; with transfers, the ratchet will halt (Takeuchi et al. 2014; Colnaghi et al. 2020). While such transfers may be rare in animal mtDNA, they are much more common in plant mtDNA (Richardson and Palmer 2007). Finally, Muller’s ratchet assumes no adaptation, as the fixation of advantageous alleles can completely stop this process (Haigh 1978; Whitlock 2000; Goyal et al. 2012). Given the large number of examples of positive selection on mtDNA (e.g. Weinreich and Rand 2000; Bazin et al. 2006; Foote et al. 2011; Morales et al. 2015; James et al. 2016), this may be the most frequently and obviously violated assumption of the ratchet model.

Our results do come with several caveats. Although *dN*/*dS* is widely used to quantify natural selection on protein-coding genes, the causes of higher or lower values can be hard to distinguish, especially in mtDNA (Adrion, White, Montooth 2016; Zwonitzer et al. 2023). The same caution should be used in interpreting *dt*/*dS*, with additional qualifications. The main one is that we have assumed that synonymous substitution rates in CDS from the same genomic compartment are a good proxy for the underlying rate of unconstrained substitution in tRNA genes. However, this may not be true. For instance, Thornlow et al. (2018) estimated that transcription-associated mutagenesis (TAM) in nuclear tRNAs caused mutation rates to be 7-10× higher at these loci. Such an effect could cause the strength of selection on nuclear tRNAs to be underestimated, but would have no direct effect on *dt*/*dS* in mtDNA tRNAs (there is no evidence for TAM in mtDNA). Overall, although there is some uncertainty associated with the measures used here, there is no evidence that they are biased for or against mtDNA loci.

Many evolutionary dynamics undoubtedly differ between mitochondrial and nuclear genomes (Ballard and Whitlock 2004), but our results suggest that researchers should be more circumspect in ascribing these differences to Muller’s ratchet. For instance, there is good evidence for coevolution between mtDNA loci and nuclear loci (Sloan et al. 2018). However, evidence for coevolution is not evidence of Muller’s ratchet, and no quantitative estimates that we know of show that there is more coevolution involving the mitochondria than in any other complex molecular machinery. Finally, and more simply, good tests for the ratchet need to distinguish this mechanism from the effects of small effective population size alone (Charlesworth et al. 1993), as the latter can lead to slightly deleterious substitutions even when there is recombination.

## Materials and methods

We obtained precomputed CDS nucleotide alignments from the UCSC Genome Browser (Perez et al. 2025). For primates, birds, and fruit flies, we used Multiz alignments of 30 mammals, Multiz alignments of 77 Vertebrates, and Multiz alignments of 27 insects, respectively. Pairwise alignments were extracted and filtered from these larger alignments using MafFilter (Dutheil et al. 2014). Orthologous genes were identified with Ensembl BioMart (Kinsella et al. 2011) for primates and birds, and with FlyBase (Thurmond et al. 2019) for fruit flies. For each orthologous CDS alignment, we estimated nonsynonymous substitutions (*dN*) and synonymous substitutions (*dS*) using the yn00 model in PAML (Yang and Nielsen 2000). We then removed CDS alignments that failed any of three filters: alignment length < 300 bp, alignment length not divisible by 3, or *dN/dS* > 2.

For tRNA genes, we used GtRNAdb annotations (Chan and Lowe 2016) on each reference genome to locate tRNA genes. Using these coordinates, we extracted and filtered pairwise tRNA gene alignments from the corresponding Multiz alignments with MafFilter as well. We estimated tRNA divergence (*dt*) for each tRNA category with baseml in PAML (Yang 2007). We removed tRNAs whose divergence was greater than 1.

All comparisons relied on additional filters applied to each pair. First, for all pairs, no genes on sex chromosomes were included. For primates, we kept 287 nuclear tRNA genes and 11690 protein-coding genes with filters mentioned above. For primate mitochondrial genes, all tRNAs were retained, but one protein-coding gene was removed because it was missing in rhesus macaque. For birds, we preserved 235 nuclear tRNA genes and 13328 protein-coding genes. Among mtDNA genes, we excluded one tRNA with extremely high divergence (*dt* = 51.3), likely due to bad alignment quality, and kept all protein-coding genes. For fruit flies, 119 nuclear tRNA genes and 8891 protein-coding genes passed filtering. All mtDNA tRNA genes were kept, whereas one protein-coding gene (*ND5*) with extremely high *dN/dS* was removed (*dN/dS* = 8.4, *dN*=0.0651, *dS*=0.0077). The final counts of all genes can be found on Supplementary Table 1.

## Acknowledgements

We thank Bryan Thornlow and Russ Corbett-Detig for sharing data. This research was supported by National Science Foundation grant DBI-2146866.

