## Supplementary Figures and Tables for "No molecular evidence for Muller’s ratchet in mitochondrial genomes"

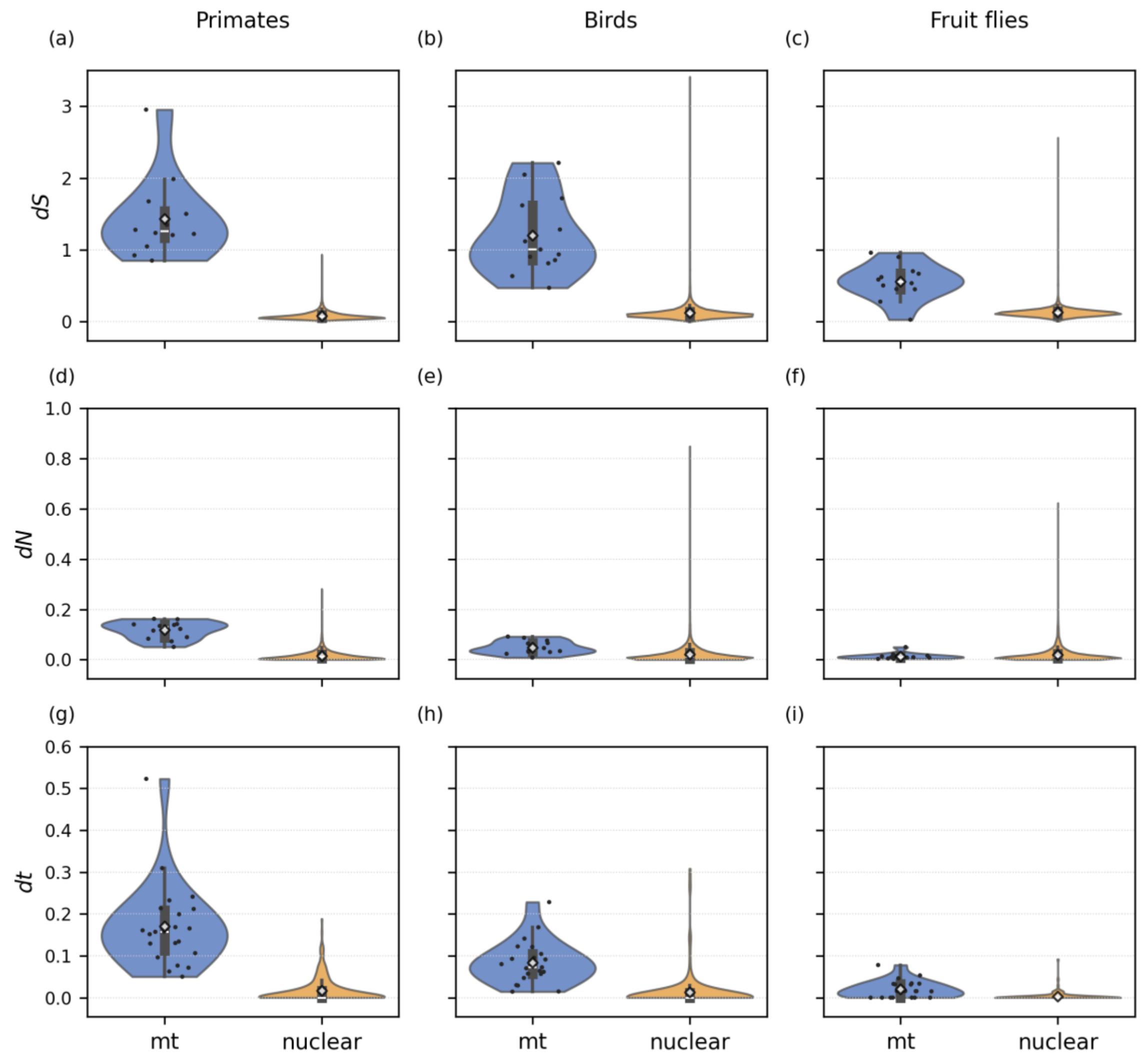


Supplementary Figure 1: Comparison between mt and nuclear genomes for (a)–(c) synonymous substitutions (*dS*), (d)–(f) nonsynonymous substitutions (*dN*), and (g)–(i) tRNA divergence (*dt*). Black dots represent data points for each mtDNA gene, and diamonds are the mean for each violin plot.

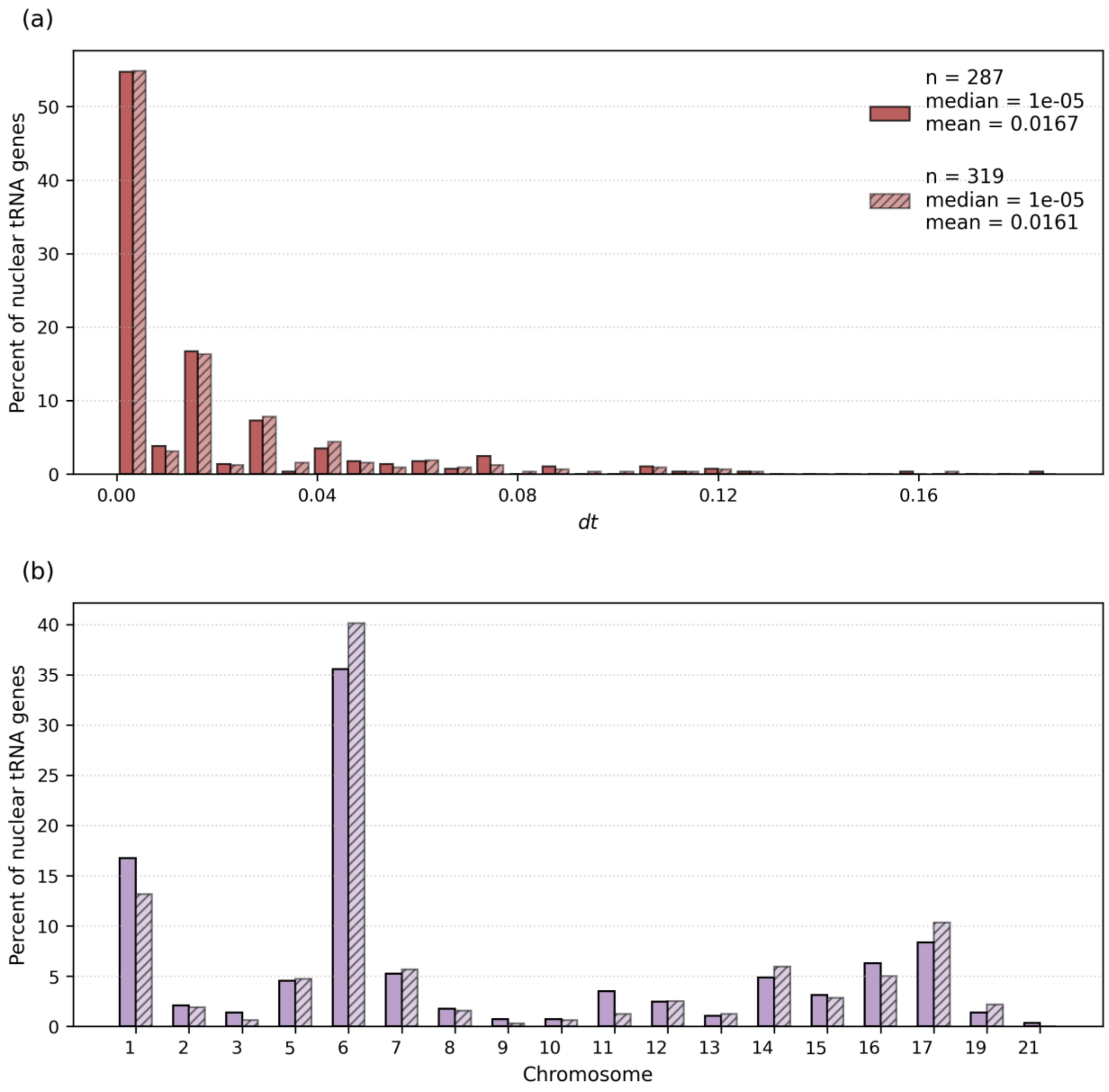


Supplementary Figure 2: Comparison of nuclear tRNA genes in primates between this study and Thornlow et al. (2018). Panels show the distributions of (a) *dt* in nuclear tRNAs, and (b) number of tRNA genes per chromosome; solid bars indicate results from this study and hatched bars show those of Thornlow et al. (2018). Similarity between datasets was assessed using two-sample Kolmogorov–Smirnov tests. Both the *dt* distributions ($D$=0.027, $p$=0.999) and chromosome assignments ($D$=0.045, $p$=0.901) were indistinguishable between datasets.

Supplementary Table 1: Number of tRNA and protein-coding genes used in each pairwise comparison.

| | Primates | | Birds | | Fruit flies | |
|---|---|---|---|---|---|---|
| | mt | nuclear | mt | nuclear | mt | nuclear |
| tRNA | 22 | 287 | 21 | 235 | 22 | 119 |
| CDS | 12 | 11690 | 13 | 13328 | 12 | 8891 |

Supplementary Table 2: Nonsynonymous substitutions and nonsynonymous-to-synonymous substitution ratio in protein-coding genes.

| | Primates | | Birds | | Fruit files | |
|---|---|---|---|---|---|---|
| | mt | nuclear | mt | nuclear | mt | nuclear |
| $dN$[a] | 0.118 | 0.0159 | 0.0486 | 0.0218 | 0.0135 | 0.02 |
| $\frac{dN}{dS}$ | 0.093 | 0.226 | 0.053 | 0.174 | 0.173 | 0.158 |

[a]average nonsynonymous per-site divergence in protein-coding genes (see Table 1 in the main text for values of $dS$)
